# ZMYM2 is essential for methylation of germline genes and active transposons in embryonic development

**DOI:** 10.1101/2022.09.13.507699

**Authors:** Adda-Lee Graham-Paquin, Deepak Saini, Jacinthe Sirois, Ishtiaque Hossain, Megan S. Katz, Qinwei Kim-Wee Zhuang, Sin Young Kwon, Yojiro Yamanaka, Guillaume Bourque, Maxime Bouchard, William A. Pastor

## Abstract

ZMYM2 is a transcriptional repressor whose role in development is largely unexplored. We found that *Zmym2^-/-^* mice show embryonic lethality by E10.5. Molecular characterization of *Zmym2^-/-^* embryos revealed two distinct defects. First, they fail to undergo DNA methylation and silencing of germline gene promoters, resulting in widespread upregulation of germline genes. Second, they fail to methylate and silence the evolutionarily youngest and most active LINE element subclasses in mice. *Zmym2^-/-^* embryos show ubiquitous overexpression of LINE-1 protein as well as aberrant expression of transposon-gene fusion transcripts. Interaction and colocalization data indicate that ZMYM2 homes to germline genes via binding to the non-canonical polycomb complex PRC1.6 and to transposons via the TRIM28 complex. In the absence of ZMYM2, hypermethylation of histone 3 lysine 4 occurs at target sites, creating a chromatin landscape unfavourable for establishment of DNA methylation. *ZMYM2^-/-^* human embryonic stem cells also show aberrant upregulation and demethylation of young LINE elements, indicating a conserved role in repression of active transposons. ZMYM2 is thus an important new factor in DNA methylation patterning in early embryonic development.

## INTRODUCTION

Methylation of the 5-position of the DNA base cytosine is the most common epigenetic modification of DNA in mammals, typically occurring at the dinucleotide CpG. DNA methylation promotes heterochromatization, and a high density of methylated CpGs at a gene promoter has a strong silencing effect^1^. DNA methylation is largely lost from the genome during pre-implantation mammalian development, then re-established globally during the periimplantation period^2^. Although methylation patterns continue to change during development, this global reprogramming in the first days of life is far more extensive than anything that occurs subsequently in somatic tissue^3^.

Most CpG-rich transcript promoters remain unmethylated even after global methylation establishment, with two categorical exceptions: germline genes and transposons. Genes selectively expressed in germ cells (germline genes) are silenced by promoter methylation and only reactivated in the developing germline, where DNA methylation is lost globally^4,5^. These genes are targeted for silencing and subsequent methylation by the non-canonical polycomb repressor complex PRC1.6, which recognizes E2F and E-Box motifs at their promoters^6,7^. Transposons are also targeted for extensive methylation. During the pre- and peri-implantation period, many transposons are silenced by the TRIM28 complex, which is recruited by hundreds of distinct KRAB-Zinc finger proteins that recognize distinct sequences present on transposons^8^. TRIM28 silences target transposons by recruiting histone deacetylases and the histone 3 lysine 9 (H3K9) methyltransferase SETDB1^8^. The TRIM28 complex is essential during development, as *Trim28^-/-^* mice show severe abnormality by embryonic day (E)5.5^9^. Additionally, TRIM28 promotes transposon DNA methylation^10,11^, and eventually DNA methylation replaces TRIM28 as the critical factor for silencing transposons^9,12,13^. Mice deficient for the *de novo* methyltransferases DNMT3A and DNMT3B show aberrant expression of germline genes and transposons along with severe growth restriction and abnormality by E8.5^13^.

The transcription factor Zinc-finger MYM-type protein 2 (ZMYM2) has been implicated in transcriptional repression in a variety of cell types^14–16^ and recruitment of ZMYM2 silences reporter constructs^14,15^. Its MYM-type Zinc fingers are reported not to bind DNA^17^, but ZMYM2 can bind the post-translational modification SUMO2/3 both via a MYM-Zinc finger and via a distinct SUMO Interaction Motif (SIM)^17,18^. Many transcription factors and chromatin-bound complexes are SUMOylated, with SUMOylation often functioning to reduce transcriptional activity^19,20^, and ectopically expressed ZMYM2 homes to regions of SUMOylation^14^. ZMYM2 may thus serve as a factor that helps mediate silencing of SUMOylated complexes on chromatin.

There is evidence that ZMYM2 mediates transcriptional repression in developmentally relevant cell types. ZMYM2 was identified as a factor important for suppressing totipotency in murine embryonic stem cells (mESCs) and zygotic depletion of *Zmym2* RNA reduces efficiency of blastocyst formation^16^. ZMYM2 was identified in a CRISPR screen as a gene that facilitates exit from pre-implantation-like “naïve” pluripotency in mESCs^21^ and has been implicated in transposon silencing in mESCs^16,22^. In subsequent CRISPR screens, ZMYM2 was found to restrict growth in human embryonic stem cells (hESCs)^23^ and its deletion results in reactivation of the silenced copy of imprinted genes in both mESCs and hESCs^24,25^. Heterozygous mutations of ZMYM2 in humans cause craniofacial abnormalities and congenital anomalies of the kidney and urinary tract (CAKUT) and *Zmym2*^+/-^ mice show elevated rates of CAKUT^15^. Nonetheless, its role and importance in mammalian embryonic development, and the molecular mechanisms underlying its role in development, remain largely unknown. To determine the developmental role of ZMYM2, we generated *Zmym2^-/-^* mice and found a striking role for ZMYM2 in facilitating the methylation of germline genes and young LINE transposons.

## RESULTS

### *Zmym2^-/-^* mouse embryos show developmental abnormality and early lethality

The *Zmym2^-^* murine allele corresponds to a mutation in the first coding exon, resulting in an early frameshift (Figure S1A). *Zmym2^-/-^* embryos were found at Mendelian ratios until E9.5, but by E10.5 resorptions were observed and no *Zmym2^-/-^* embryos were recovered, demonstrating embryonic lethality (Figure 1A). A variable *Zmym2^-/-^* phenotype was observed at E9.5, with most embryos showing gross phenotypic abnormality, including a failure to undergo turning of the embryonic trunk (Figure 1B,C). A statistically significant reduction in length, size and somite count was also observed (Figure 1D, Figure S1B,C).

**Figure 1.**
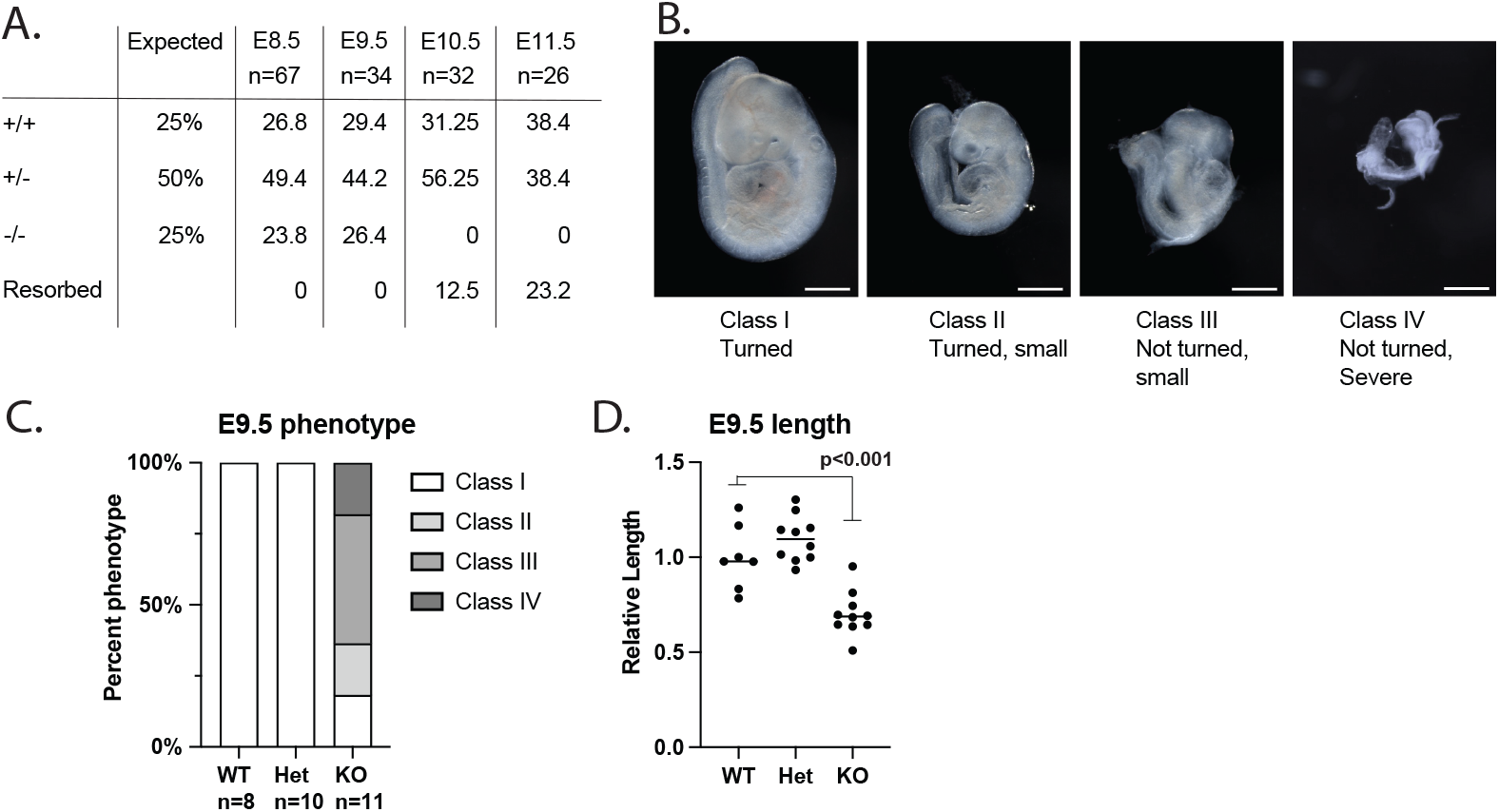
*Zmym2^-/-^* embryos show lethality by E10.5 and variable developmental delays at E9.5. **A.** Embryonic viability by genotype from embryonic day E8.5 to E11.5 compared to the expected Mendelian ratios from *Zmym2^+/-^; Zmym2^+/-^* crosses. Each embryonic age includes pooled data from 3+ litters. **B.** Images taken of E9.5 *Zmym2^-/-^* embryos representing four different classifications of phenotype severity. Scale bars= 500μm. **C.** Distribution of E9.5 phenotype classes by genotype. **D.** Relative length of embryos (crown to tail) normalized by litter as a ratio to average *Zmym2^+/+^* length. Significance from one sided t-test is indicated.

Most E8.5 *Zmym2^-/-^* embryos appeared normal (Figure S1D), so the molecular phenotype of *Zmym2^-/-^* was analyzed using E8.5 embryos with no visible abnormalities.

### Zmym2 suppresses expression of germline genes and young LINE elements

To determine the transcriptional changes that occur in *Zmym2^-/-^* embryos prior to embryonic lethality, we performed RNA-sequencing of six *Zmym2^+/+^* and six *Zmym2^-/-^* whole E8.5 embryos, three male and three female of each genotype (Tables S1, S2). Upon the loss of ZMYM2, knockout embryos exhibited an upregulation of 165 genes (fold change ≥ 4, q-value < 0.05) (Figure 2A), and downregulation of only three genes, consistent with a repressive role for ZMYM2. GO term analysis showed that the upregulated genes were enriched for terms related to germ cell development (Figure 2B). Upregulation of select genes was confirmed through RT-qPCR of additional *Zmym2^-/-^* and *Zmym2^+/+^* E8.5 embryos (Figure S1E, Table S3).

**Figure 2.**
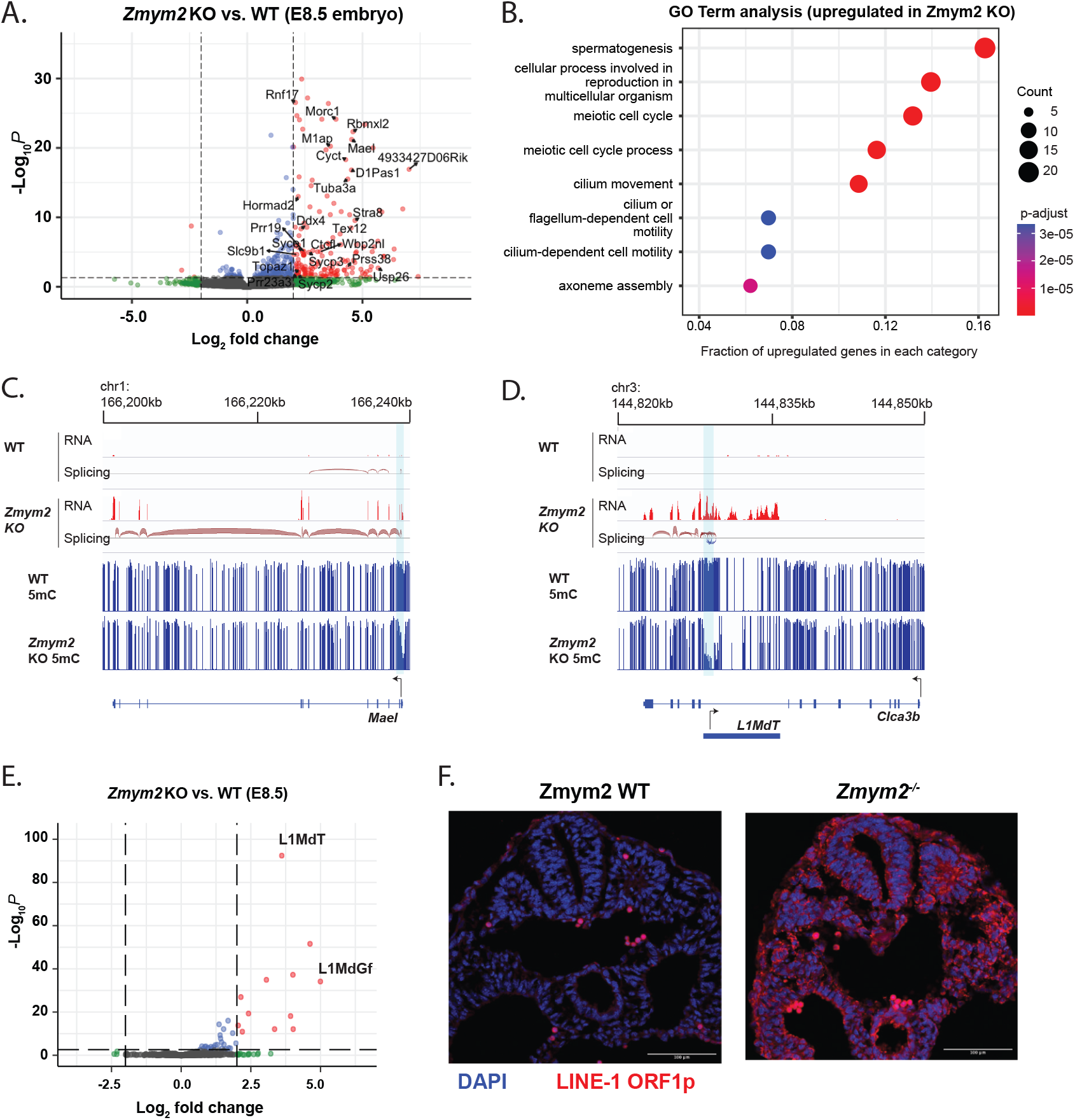
Germline genes and young LINE elements are upregulated in *Zmym2^-/-^* embryos. **A.** Volcano plot of differentially expressed genes comparing *Zmym2^+/+^* and *Zmym2^-/-^* E8.5 embryos. Significant differentially expressed genes (fold ≥ 4, q-value < 0.05) are expressed in red. Germline genes are labeled. The upregulated gene *Pla2g1b* is off-axis and not plotted. **B.** Gene ontology (GO) analysis of significantly upregulated genes (fold change ≥ 4, q-value < 0.05). **C.** Transcription and DNA methylation over the germline gene *Mael*. Note upregulation in *Zmym2^-/-^* and promoter hypomethylation. A hypomethylated DMR is indicated in light blue. **D.** Transcription and DNA methylation over the gene *Clca3b*. Note splicing from the L1MdT element to the *Clca3b* transcript, as well as hypomethylation of the L1MdT promoter. A hypomethylated DMR is indicated in light blue. **E.**Volcano plot of differentially expressed transposable elements comparing *Zmym2^+/+^* and *Zmym2^-/-^* E8.5 embryos. Significant differentially expressed transposons (fold ≥ 4, q-value < 0.05) are coloured red. **F.** Immunofluorescence staining of LINE-1 ORF1p (red) and DAPI (blue) in E9.5 *Zmym2^+/+^* and *Zmym2^-/-^* embryonic tissue. Note that the few, bright red cells in *Zmym2^+/+^* are autofluorescent red blood cells. Scale bars= 500μm.

We observed two classes of upregulated genes in *Zmym2^-/-^* embryos. Ninety-five genes showed upregulation of transcription from an annotated promoter (Figure 2C). By contrast, 60 genes showed clear evidence of splicing from an internal or upstream transposon (Figure 2D, annotations in Table S4). In 46 of the 60 fusion transcripts, the transposon in question was a LINE element of the L1MdT subclass. Both sense and antisense transcription from LINE promoters seeded transcription of fusion genes (Figure 2D, Figure S2A-C). Interestingly, published data^26^ indicates that some of these LINE-gene fusion transcripts are expressed at relatively high levels in early development and suppressed between E6.5 and E8.5 in wild-type mice (Figure S2D-F).

We analyzed transposon expression in *Zmym2^-/-^* using the TEtranscript pipeline^27^ and observed strong upregulation of L1MdT and the less abundant L1MdGf subclasses (Figure 2E). In any given species, the youngest transposons classes are typically the most active and capable of transposition, and it is notable that L1MdT and L1MdGf are the youngest and third youngest LINEs in *Mus musculus* respectively^28^. Immunofluorescence staining of cross-sectional slices of E8.5 and E9.5 *Zmym2^-/-^* embryos showed high expression of the LINE-1 ORF1 protein (L1ORF1p) in all cells, in striking contrast with *Zmym2^+/+^* controls (Figure 2F, Figure S2G). L1ORF1p is also highly expressed in *Zmym2^-/-^* trophoblast giant cells, indicating that ZMYM2 is also important for suppressing transposons in the placental lineage (Figure S2H).

### Zmym2 is required for DNA methylation of germline gene promoters and LINE elements

As germline genes and transposons are known to be silenced by DNA methylation in the somatic lineage, we conducted whole genome bisulfite sequencing (WGBS) of nine *Zmym2^+/+^* and nine *Zmym2^-/-^* embryos. We targeted>30-fold coverage for male and female, *Zmym2^+/+^* and *Zmym2^-/-^* (Table S1). Two embryos, one of each genotype and originating from the same litter, were excluded from subsequent analysis because they showed abnormally low levels of global DNA methylation for E8.5 (Figure 3A).

**Figure 3.**
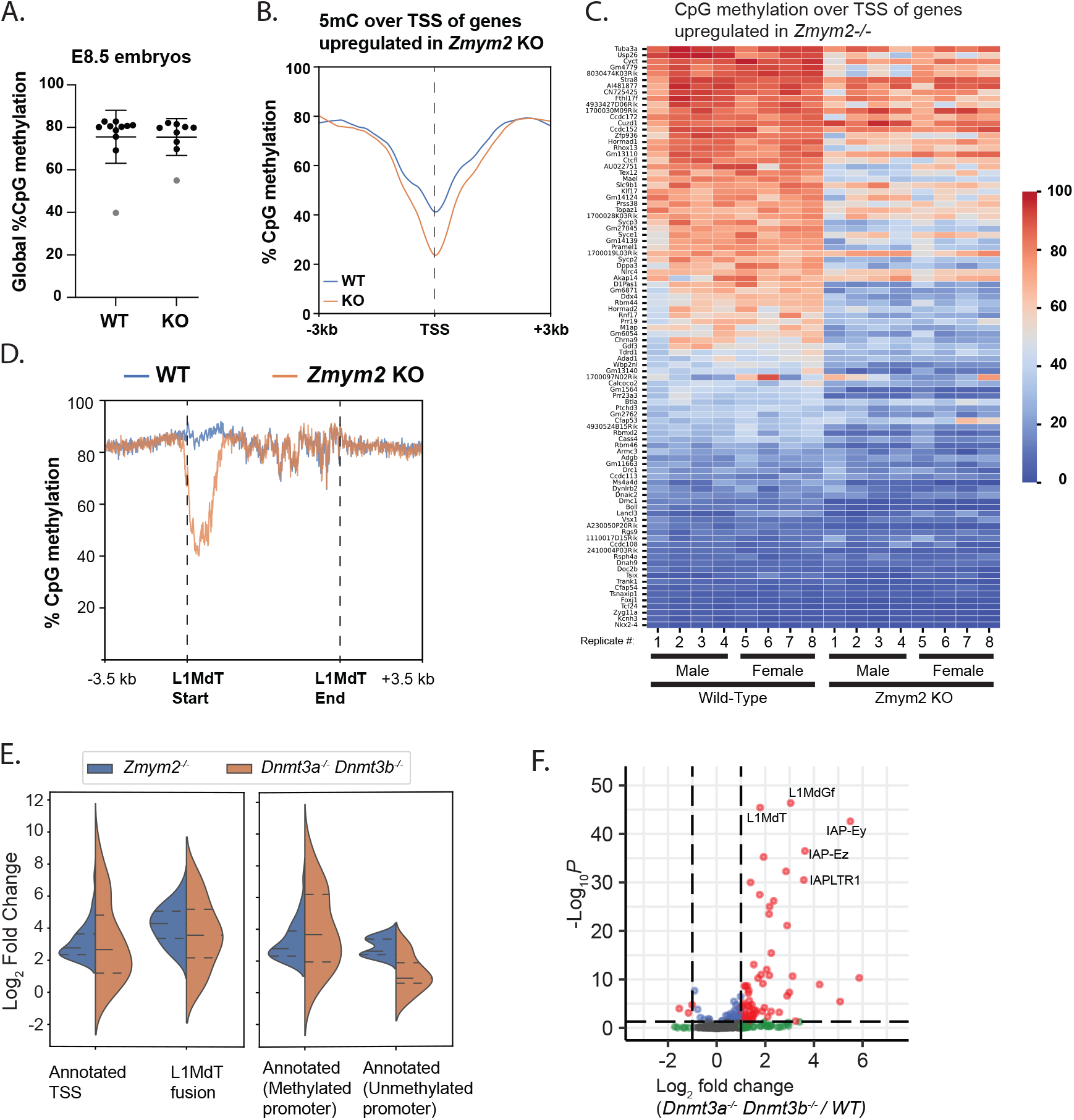
Impaired DNA methylation in *Zmym2^-/-^* over germline genes and LINE elements. **A.** Swarm plot of global CpG methylation levels of individual *Zmym2^+/+^* and *Zmym2^-/-^* E8.5 embryos. Two samples coloured gray were excluded from subsequent analysis **B.** Metaplot of CpG methylation over TSS of 95 upregulated genes in *Zmym2^-/-^* embryos expressed from annotated TSS. **C.** Heatmap of DNA methylation of the TSS within 500bp of the transcription start site of each of upregulated gene is indicated for each E8.5 embryo. Transposon-fusion genes and the Y-chromosomal gene *Uba1y* are excluded. **D.** Metaplot of DNA methylation over all full-length (>6kb) L1MdT elements in the genome. **E.** Expression of each gene in the given category is shown (KO/WT) in *Zmym2^-/-^* and *Dnmt3a^-/-^ Dnmt3b^-/-^*. Note that most genes upregulated in *Zmym2^-/-^* show elevated expression in *Dnmt* double knockout. **F.** Volcano plot of differentially expressed transposable elements comparing *Dnmt3a^-/-^ Dnmt3b^-/-^* and control E8.5 embryos.

We observed no defect in global DNA methylation establishment in *Zmym2^-/-^* embryos (Figure 3A). However, we observed a substantial reduction in DNA methylation over the promoters of the 95 upregulated genes expressed from annotated TSS (Figure 2C, Figure 3B,C). Hypomethylation was also observed globally over the promoters of full length L1MdT elements (Figure 2D, Figure 3D, Figure S2A-C). We compared gene expression changes in the *Zmym2^-/-^* embryos to published data^13^ of E8.5 *Dnmt3a^-/-^/3b^-/-^* mice, in which all *de novo* methylation is impaired. Genes upregulated in *Zmym2^-/-^* and expressed from annotated TSS showed a globally similar degree of upregulation in *Dnmt3a^-/-^/3b^-/-^* embryos, consistent with their normally being silenced by DNA methylation (Figure 3E). When this subgroup was subdivided into genes with methylated (>20% CpG methylation at TSS) and unmethylated promoters, higher upregulation in *Dnmt3a^-/-^/3b^-/-^* was observed in the methylated set (Figure 3E). L1MdT-fusion transcripts were also upregulated in *Dnmt3a^-/-^/3b^-/-^*, although not to the same extent as in *Zmym2^-/-^* (Figure 3E). Likewise, more overall transposon classes were upregulated in *Dnmt3a^-/-^/3b^-/-^* than *Zmym2^-/-^* but upregulation of L1MdT and L1MdGf elements was somewhat weaker, suggesting that ZMYM2 may silence young LINEs by both methylation-dependent and -independent mechanisms (Figure 3F).

Further consistent with a role for ZMYM2 in promoting DNA methylation, ZMYM2 expression peaks during post-implantation development in both mouse^26^ and primate^29^, coincident with expression of *de novo* methyltransferases (Figure S3A,B). As with L1MdT-fusion transcripts, ZMYM2 target genes generally show downregulation between E6.5 and E8.5, consistent with their being silenced and methylated during this time (Figure S3C).

We then identified differentially methylated regions (DMRs) using a 1,000bp tiling approach. Three thousand three hundred forty such regions were hypomethylated in *Zmym2^-/-^*, and only 224 regions were hypermethylated (Table S5). 40.9% of these hypomethylated DMRs overlap with L1MdT elements, compared with 2.6% overlap expected by chance (Figure S3D). Other LINE elements, most notably L1MdGf, also overlapped with DMRs more than expected by chance (Figure S3E-G). This approach confirms ZMYM2’s role in directing DNA methylation.

We then examined other types of loci regulated by DNA methylation. Methylation of gene promoters on the X-chromosome proceeded normally in female *Zmym2^-/-^* mice, indicating that ZMYM2 was not essential for methylation after X-inactivation (Figure S3H). Canonical imprinted loci, which inherit methylation from parental gametes, do not show hypomethylation (Figure S3I), consistent with ZMYM2’s role being establishment rather than maintenance of methylation. Intriguingly though, we observed hypomethylation and transcription of the transient imprint *Liz* (Long isoform of *Zdbf2*) (Figure S3J), a locus which is imprinted during preimplantation development but becomes biallelically methylated and fully silenced during the peri-implantation period^30^.

### ZMYM2 methylates young LINE retrotransposons in human ESCs

We used published RNA-seq and reduced representation bisulfite-seq (RRBS) data from *ZMYM2^-/-^* hESCs^23,24^ to determine if dysregulation of transposons was observed. Indeed, we observed highly significant upregulation of the L1Hs and L1PA2 sub-families, the two youngest LINE element families in humans (Figure 4A). This was not a result of the modest shift toward a pre-implantation “naïve” gene expression signature observed in *ZMYM2^-/-^* hESCs^23^ because naïve cells do not show substantial increases in L1Hs and L1PA2^31^ (Figure S4A). Individual L1Hs elements are too similar to each other to assign individual RRBS reads to individual elements efficiently, but mapping to a common consensus sequence showed that loss of *ZMYM2* or its known protein interactor *ATF7IP*^15,22^ results in hypomethylation of the L1Hs promoter region in hESCs (Figure 4B, Figure S4B,C). Strong hypomethylation of L1PA2 elements is also observed in *ZMYM2^-/-^* and *ATF7IP^-/-^* (Figure 4C, Figure S4D,E). Thus, ZMYM2’s role in methylating young transposons is conserved in humans.

**Figure 4.**
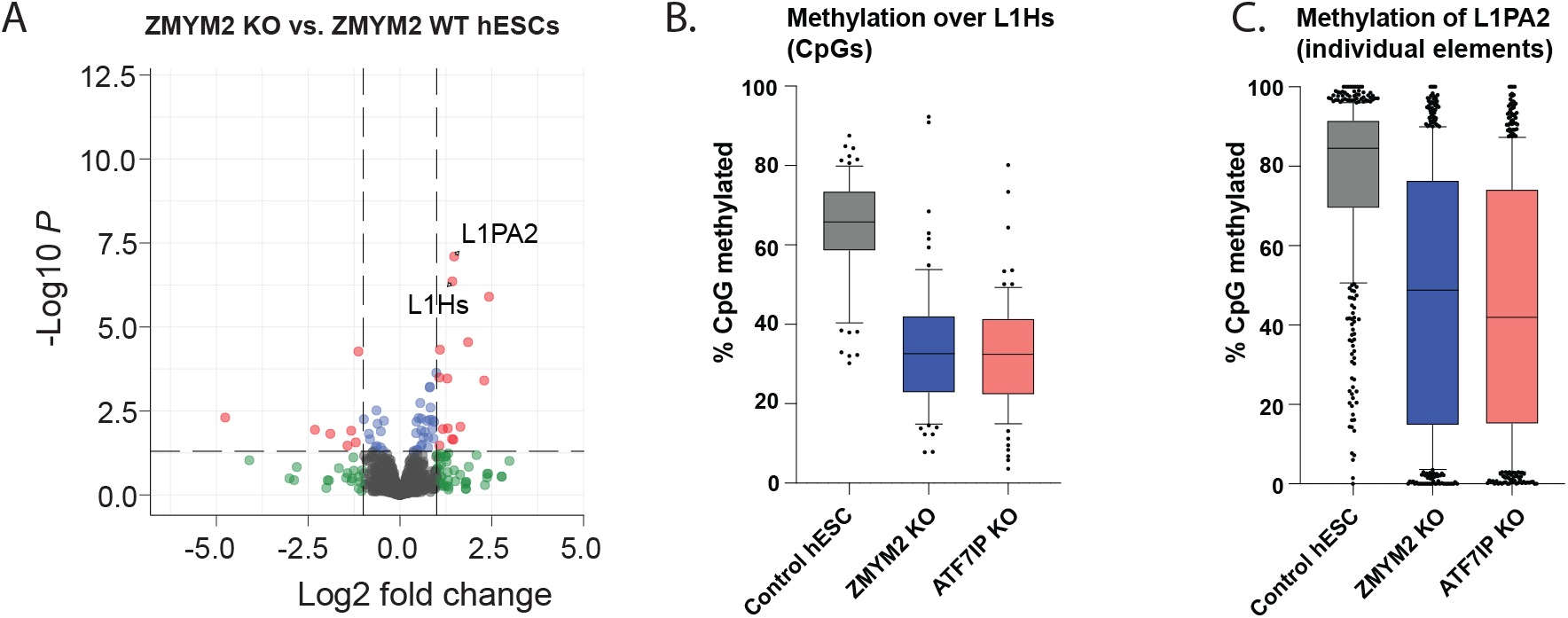
Upregulation and hypomethylation of young LINE elements in *ZMYM2^-/-^* human embryonic stem cells. **A.** Volcano plot of differentially expressed transposons comparing *ZMYM2^+/+^* and *ZMYM2^-/-^* hESCs. Significant differentially expressed transposons (fold ≥ 2, q-value < 0.05) are coloured red. **B.** Boxplot of CpG methylation across 1Kb start regions of full length L1Hs transposable elements. Data was mapped to an L1Hs consensus element, and individual CpG sites are represented as dots. **C.** Boxplot of average CpG methylation across L1PA2 individual transposable elements. Each element is represented as a dot.

### ZMYM2 silences target loci in embryonic stem cells and embryoid bodies

We generated *Zmym2^-/-^* mESC lines on a V6.5 (mixed C57B6/129) background (Figure S5A-B). *Zmym2^-/-^* mESCs could be differentiated to embryoid bodies (EBs), showing similar upregulation of differentiation markers as *Zmym2^+/+^* mESCs (Figure 5A). By day 6 of differentiation reduced EB size and viability was observed, mimicking the lethality observed in mice (Figure S5C).

**Figure 5.**
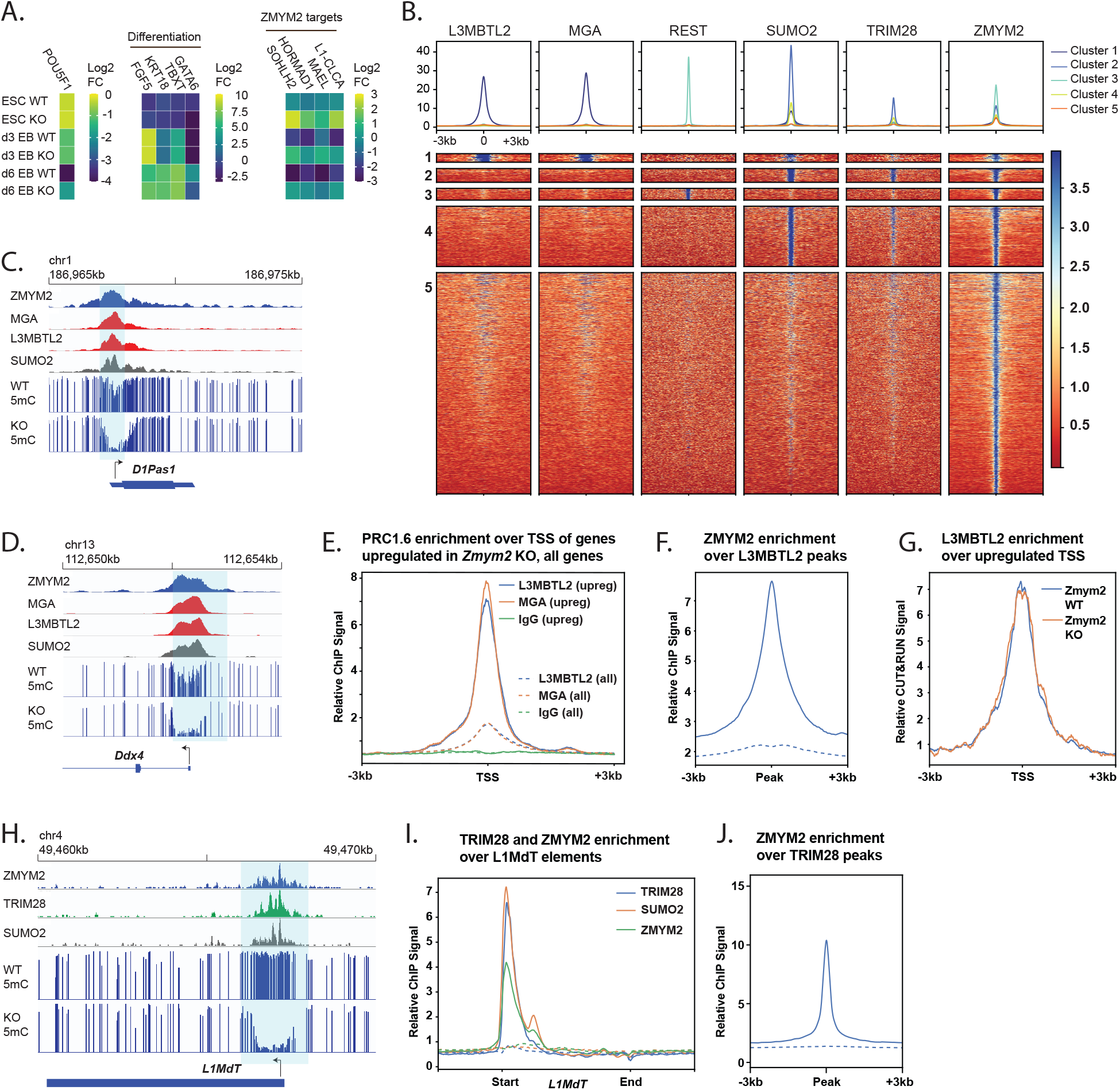
ZMYM2 silences PRC1.6 and TRIM28 targets and promotes their methylation. **A.** Relative expression of the pluripotency factor *Pou5f1*, differentiation genes, and ZMYM2-target genes in three *Zmym2^-/-^* lines as compared with three control lines, during two separate embryoid body differentiation experiments. Average results from these n=6 replicates total are shown. **B.** Heatmaps of ZMYM2, L3MBTL2, MGA, REST, SUMO2, and TRIM28 enrichment over ZMYM2 ChIP-seq peak summits and the flanking 3Kb regions (n = 21,549 peaks). Peaks were partitioned into five clusters using a k-means algorithm. Peak numbers: Cluster 1, 290 peaks; Cluste 2, 931 peaks; Cluster 3, 796 peaks; Cluster 4, 4080 peaks; Cluster 5, 14969 peaks. Cluster 1 peaks are shown at twice the width of other peaks to enhance visibility. **C.,D.,** ChIP-seq and DNA methylation data over the *D1Pas* (**C**) and *Ddx4* (**D**) loci. Note colocalization of ZMYM2 with PRC1.6 components and hypomethylation in *Zmym2^-/-^*. **E.** A Metaplot of L3MBTL2, MGA and IgG negative control ChIP-seq signals over the annotated transcription start site of upregulated genes in *Zmym2^-/-^* E8.5 embryos (solid lines) with transposon-fusions excluded and the annotated transcription start site of all genes (dashed lines). **F.** Metaplot of ZMYM2 ChIP-seq signal over L3MBTL2 peaks. Dashed line indicates ChIP input. **G.** Metaplot of L3MBTL2 CUT&RUN ChIP-seq signal over the annotated TSS of upregulated genes in both *Zmym2^+/+^* and *Zmym2^-/-^* mESCs. **H.** ChIP-seq and DNA methylation data over a representative LINE element. Note overlap of TRIM28 and ZMYM2 peaks and hypomethylation over the peak site. **I.** Metaplot of ZMYM2, TRIM28, SUMO2 ChIP-seq over all full length L1MdT elements. Dashed line indicates ChIP input. **J.** Metaplot of ZMYM2 over all uniquely mapped TRIM28 peaks. Dashed line indicates ChIP input.

*Zmym2^-/-^* mESCs showed upregulation of germline genes *(Mael, Hormad1, Sohlh2)* and a transposon fusion *(L1MdT-Clca3b)* relative to control *Zmym2^+/+^* mESC lines (Figure 5A). Upon differentiation to EBs, expression of the ZMYM2-target transcripts dropped substantially in *Zmym2^+/+^* mESCs but to a lesser degree in *Zmym2^-/-^*, accentuating the relative upregulation (Figure 5A), akin to the phenomenon by which ZMYM2 targets are suppressed in *Zmym2^+/+^* but not *Zmym2^-/-^* mouse embryo development (Figure S2F, S3C). Similar results were observed from a *Zmym2^-/-^* CCE mESC line from a pure 129/Sv genetic background^16^ (Figure S5D). Thus, *Zmym2^-/-^* mESCs and EBs recapitulate dysregulation of germline genes and transposons and can be used as an *in vitro* model with which to study ZMYM2 targeting and activity.

### ZMYM2 silences genes and transposons downstream of PRC1.6 and TRIM28 respectively

ZMYM2 lacks capacity for sequence-specific DNA binding but can bind SUMOylated transcription factors or complexes on chromatin and act as a corepressor^22,32^. As discussed above, PRC1.6 is essential for silencing of germline genes in early embryonic development while TRIM28 silences transposons. Also, we noted that a previously reported ZMYM2-motif^16^ is almost identical to the binding motif of the transcription factor REST. We thus performed cluster analysis of all high-enrichment ZMYM2 ChIP-seq peaks, incorporating published mESC ChIP-seq data for ZMYM2^16^, PRC1.6 components (MGA and L3MBTL2^7^), TRIM28^33^, REST^34^ and SUMO2^33^. We observed strongly distinct PRC1.6^+^, TRIM28^+^ and REST^+^ clusters, indicating that ZMYM2 homes to each of complexes independently (Figure 5B).

ZMYM2 is known to interact with the PRC1.6 complex^15,16^, which is heavily SUMOylated in mESCs^35^. ZMYM2 tightly colocalizes with PRC1.6 at promoters of germline genes in mESCs (Figure 5C,D, S5E,F) and PRC1.6 is strongly enriched at genes upregulated in *Zmym2^-/-^* mouse embryos relative to other genes (Figure 5E). ZMYM2 is globally enriched over PRC1.6 ChIP-seq peaks (Figure 5F), with virtually all PRC1.6 sites showing enriched binding of ZMYM2 (Figure S5G). We conducted CUT&RUN to detect distribution of the PRC1.6 component L3MBTL2 in control and *Zmym2^-/-^* mESCs and found no difference in binding to genes upregulated in *Zmym2^-/-^* embryos (Figure 5G). Likewise, there is no loss of the histone modification H2AK119Ub (deposited by the RING1 component of PRC1.6) in *Zmym2^-/-^* mESCs or day 3 EBs (Figure S5H,I). Thus, ZMYM2 is not essential for PRC1.6 binding or activity but is nonetheless essential as a corepressor to silence some PRC1.6 target genes.

The TRIM28 complex is heavily SUMOylated in mESCs^35^, and ZMYM2 could also home with the TRIM28 complex via their known mutual interactor ATF7IP^22^. ZMYM2 shows colocalization with TRIM28 at L1MdT elements, and ZMYM2 is strongly enriched over TRIM28 binding sites genomewide (Figure 5H-J, Figure S5G,J,K). As with PRC1.6, ZMYM2 homes to virtually all TRIM28 targets and is essential for silencing of a distinct subset of them.

How ZMYM2 homes to REST bindings sites is unclear. ZMYM2 and REST both bind the CoREST complex but in a mutually exclusive manner^17^ and REST is not known to be SUMOylated. Nonetheless, ZMYM2 is very strongly enriched at REST sites (Figure 5B, Figure S5L,M). We do not observe hypomethylation of REST sites (Figure S5N) or upregulation of REST target genes^36^, so the biological significance of this interaction remains unclear.

### Loss of ZMYM2 results in H3K4 hypermethylation at target loci

The histone modifications H3K4me2 and H3K4me3 are ubiquitous at sites of transcriptional initiation in mammals^37,38^. The *de novo* methyltransferases DNMT3A, DNMT3B and DNMT3L in turn bind to the H3 N-terminus via their ADD domains, but this binding is strongly antagonized by the presence of the modification H3K4me2 or H3K4me3^39–41^. H3K4me2/3-marked chromatin thus escapes *de novo* DNA methylation during development^42–44^, and actively transcribed gene promoters remain unmethylated in the post-implantation embryo^45–47^. PRC1.6 and TRIM28 both silence target loci prior to deposition of methylation^9,48^. We therefore theorized that ZMYM2 may promote DNA methylation by causing transcriptional silencing and loss of H3K4me2/3 at target sites.

We performed ChIP-seq of H3K4me2 and H3K4me3 in *Zmym2^-/-^* mESCs and day 3 EBs as well as control *Zmym2^+/+^* lines and normalized each replicate to control for ChIP quality (Figures S6A-D). We observe elevated levels of H3K4me2 and H3K4me3 in *Zmym2^-/-^* mESCs over the hypomethylated DMRs found in E8.5 *Zmym2^-/-^* mice (Figure 6A,B). H3K4 hypermethylation was also observed at the promoters of full-length L1MdT elements and the genes upregulated in *Zmym2^-/-^* mice (Figure 6C-H, Figure S6E-G). These trends are accentuated further upon differentiation to EBs, as H3K4 levels over ZMYM2-regulated loci drop substantially in *Zmym2^+/+^* cells but much less so in *Zmym2^-/-^* cells (Figure 6, Figure S6). Hence, ZMYM2 functions to reduce H3K4me2/3 at target genes and transposons and thus creates a chromatin environment conducive to *de novo* methylation.

**Figure 6.**
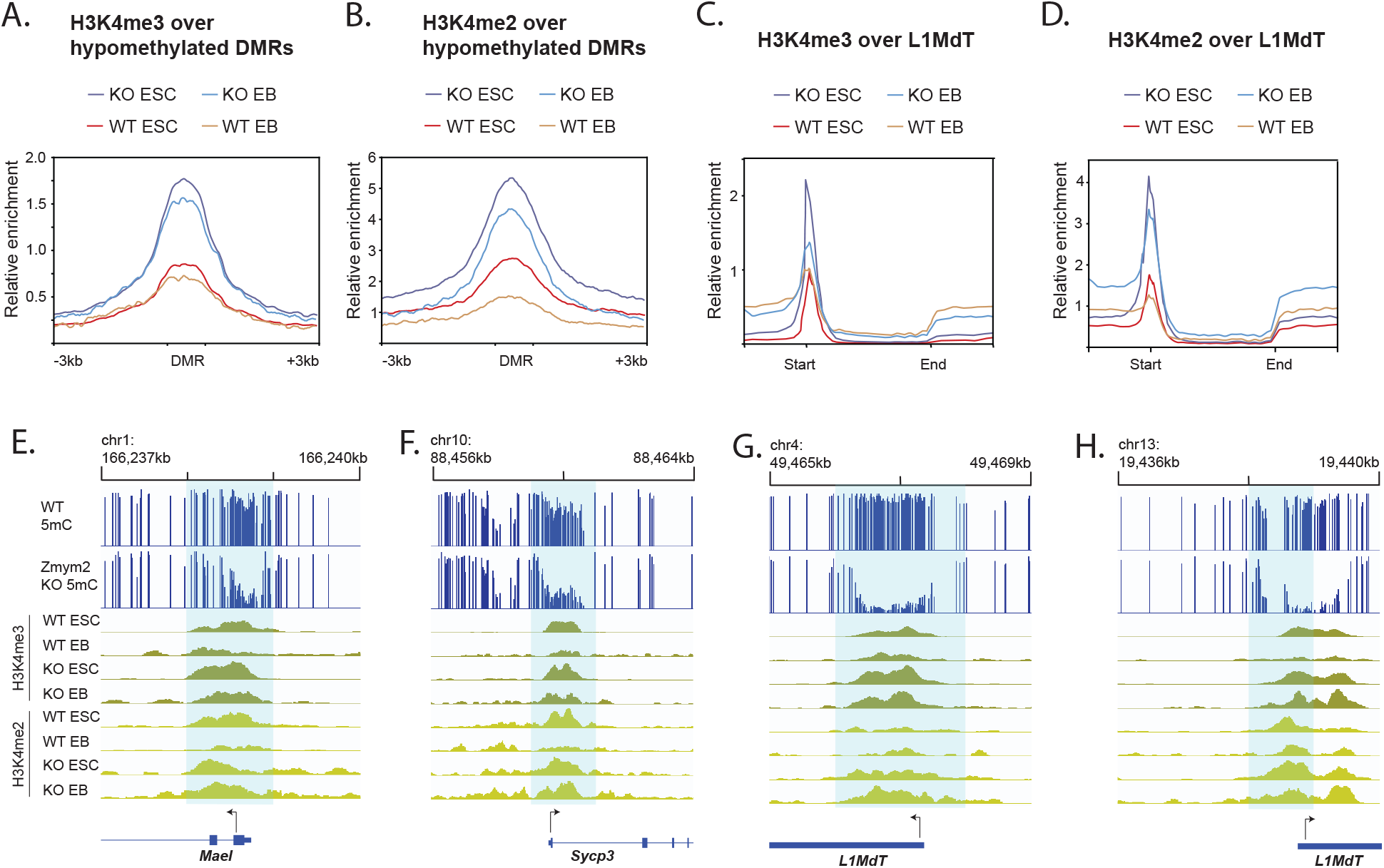
Aberrant H3K4 hypermethylation at ZMYM2-target sites in *Zmym2^-/-^* mESCs and EBs. **A.** Metaplot of H3K4me3 over DMRs hypomethylated in *Zmym2^-/-^* embryos in *Zmym2^-/-^* and control ESCs and EBs. **B.** Metaplot of H3K4me2 over hypomethylated DMRs. **C.** Metaplot of H3K4me3 over full length L1MdT elements. **D.** Metaplot of H3K4me2 over full length L1MdT elements. For **A-D**, averaged data for two *Zmym2^-/-^* and two control mESC lines are shown. To normalize for ChIP quality, enrichment is normalized based on peak height for all annotated genes. **E – H.** Examples of increased H3K4me2/3 in *Zmym2^-/-^* mESCs over hypomethylated DMRs including genes (**E,F**) and transposons (**G,H**).

## DISCUSSION

ZMYM2 is clearly essential for silencing and DNA methylation of germline genes and young transposons in development. A combined model of its activity runs as follows. ZMYM2 binds to PRC1.6 and TRIM28 sites, where it functions as a corepressor. By suppressing transcription, it reduces levels of H3K4 methylation, facilitating DNA methylation upon implantation and establishing stable silencing (Figure 7).

**Figure 7.**
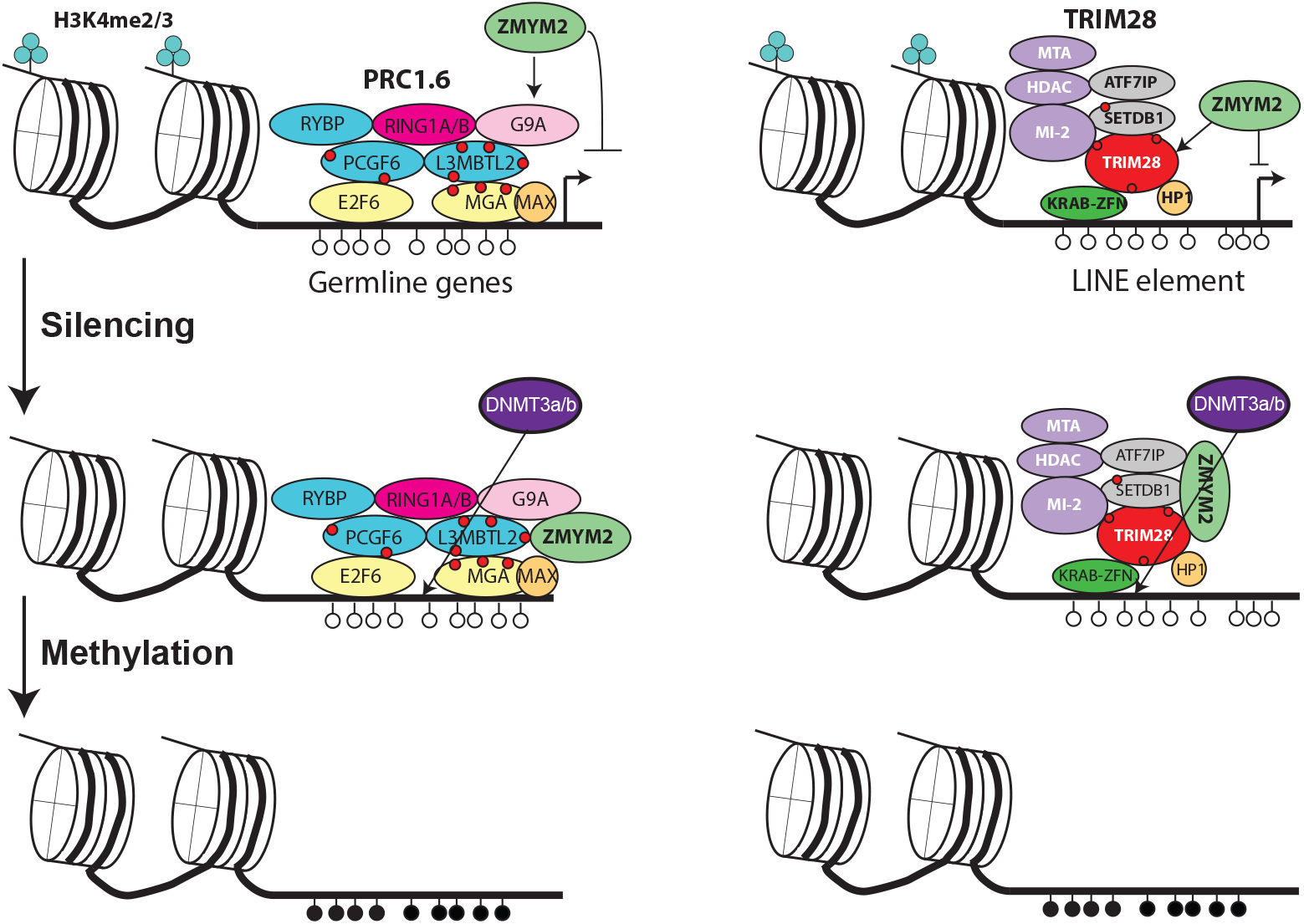
Proposed mechanism for ZMYM2-mediated DNA methylation. ZMYM2 is recruited to PRC1.6 and TRIM28 binding sites respectively via binding to SUMOylation sites (indicated as red circles) or interaction with complex components such as ATF7IP. ZMYM2 mediates silencing of target loci, resulting in loss of H3K4 methylation and acquisition of DNA methylation.

PRC1.6 and TRIM28 are heavily SUMOylated in mESCs^35^ and ZMYM2 could home to them via these SUMO2 marks (Figure 5). At the same time, we cannot rule out SUMO-independent recruitment of ZMYM2 to these complexes. ZMYM2 interacts with the Fibronectin III domain of ATF7IP in an apparently SUMO-independent manner, and ATF7IP has been reported to interact with both TRIM28 and the PRC1.6 component MGA^22^. It is striking that loss of either ATF7IP or ZMYM2 results in hypomethylation of young LINE elements in hESCs (Figure 4B,C, Figure S4B-E), and both genes have been hits in CRISPR screens for factors that maintain imprint methylation in stem cells^24,25^. One possibility is that ATF7IP’s primary function at these loci is to recruit ZMYM2 to such elements. However, ATF7IP performs other function such as promoting nuclear localization of the H3K9 methyltransferase SETDB1^49^, so the similarity of the *ATF7IP^-/-^* and *ZMYM2^-/-^* phenotypes could reflect genuine interaction or just that both proteins are important for silencing.

How ZMYM2 mediates silencing at germline genes and transposons is not certain, but there is extensive evidence that ZMYM2 binds to the CoREST complex^15–17,50,51^, which includes the histone deacetylases HDAC1 and HDAC2 and the histone demethylase LSD1^50^. Loss of ZMYM2 in HeLa cells or mESCs, is reported to result in reduced global association of CoREST with chromatin^16,17^. ZMYM2 may thus silence targets by recruiting CoREST and its associated histone deacetylases. LSD1 recruitment could also cause H3K4me1 and H3K4me2 demethylation^52^, which would promote silencing and facilitate downstream DNA methylation.

The *Zmym2^-/-^* mice show embryonic lethality amidst aberrant upregulation of LINE protein. It is difficult to determine what precisely halts the development of the *Zmym2^-/-^* embryos, but it is well established that LINE elements can induce cellular lethality, both by inducing anti-viral responses and by the endonuclease activity of L1ORF2p^53^. The phenomenon of LINE-gene fusion transcripts warrants further note. There are numerous examples of LTR-retrotransposons, most frequently “solo LTRs” detached from larger transposons, serving as alternative promoters or enhancers for protein-coding transcripts^54,55^. LINE elements integrate into the genome in a 3’ to 5’ manner, so a promoter is only present if complete integration has occurred. Thus, “orphan” LINE promoters are expected to be a rarer phenomenon. Nonetheless, LINE-gene fusions apparently occur, both as a result of bidirectional and forward transcription, and are expressed at significant levels in wild-type pre-implantation embryos. We do not know how many of the 46 LINE-transcript fusions we observe code for functional protein or whether they have any biological function, but it is intriguing that L1MdT expression during pre-implantation murine development is essential for developmental progression^56,57^.

It is important to consider the phenotypes we do not see in *Zmym2^-/-^* but might have expected based on literature. We do not see pre-implantation developmental arrest^16^, but this can be explained by maternal deposition of intact *Zmym2* RNA in the oocytes of *Zmym2^+/-^* mothers. A number of published observations in embryonic stem cells were not observed in the mice. We also do not observe failure to exit pluripotency^21,23^, with E8.5 *Zmym2^-/-^* having undergone gastrulation normally. Embryonic stem cells adapted to continuous pluripotent culture may be more resistant to differentiation than developing mice. We also do not observe loss of canonical imprints in *Zmym2^-/-^* embryos^24,25^ (Figure S3I). It may be that the dynamic state of mESCs, in which both *de novo* methylation and demethylation occur in continuous culture, does not reflect methylation dynamics in the rapidly developing mouse embryo. Alternatively, ZMYM2 may be essential to maintain imprints during the pre-implantation period when DNA methylation is lost, and maternally-inherited *Zmym2* masks this phenotype in the *Zmym2^-/-^* mice.

Unlike the *Zmym2^-/-^* mice, we also do not observe demethylation of germline genes in *ZMYM2^-/-^* hESCs (data not shown). While this could reflect species difference, it is important to note here that conventionally cultured primed hESCs correspond to a developmentally advanced state that features high global DNA methylation, including over germline genes^31^. Even if ZMYM2 is critical for DNA methylation establishment over germline genes in development, its loss will not necessarily cause loss of existing DNA methylation. Young LINE elements by contrast are targets of continuous TET activity in hESCs and thus depend on continual *de novo* methyltransferase activity to remain methylated^58^, potentially explaining how they become demethylated upon loss of ZMYM2.

ZMYM2 shows co-association with a variety of chromatin-bound complexes. Zygotic depletion of *Zmym2* RNA blocks blastocyst formation^16^. The *Zmym2^-/-^* mouse we have generated, which is to our knowledge the first report of a ZMYM-family transcription factor knockout in literature, shows lethality and severe epigenetic abnormality. The cranial, cardiac, musculoskeletal, CAKUT and possible infertility phenotype caused by ZMYM2 heterozygosity in humans suggests extensive further roles in development^15^. Mutations of ZMYM3 has been implicated in mental retardation^59,60^, and all six ZMYM-family transcription factors are widely expressed in human tissues^61^. The role of ZMYM-family transcription factors as cofactors for repression may be widespread and underappreciated.

## FIGURE CAPTIONS

**Figure S1.**
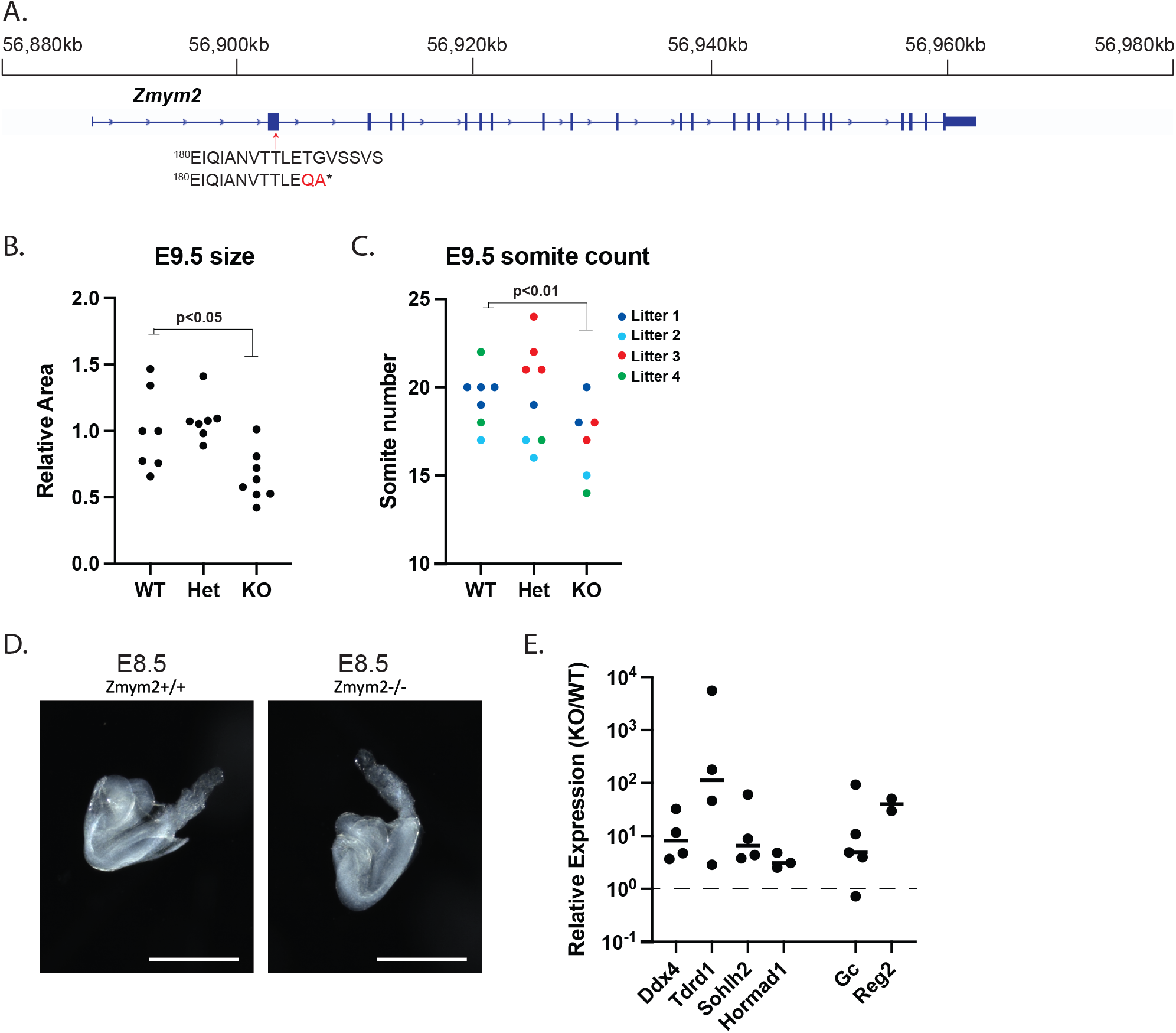
Reduced size and developmental delays in *Zmym2^-/-^* mice. **A.** Schematic representation of point mutation and truncation point in *Zmym2^-^* allele. **B.** Relative size of embryos measured as 2D area in embryos in sagittal position normalized by litter as a ratio to average *Zmym2^+/+^* area. p-value calculated on basis of one-tailed t-test. **C.** Somite counts of E9.5 embryos per genotype. Each litter is represented in a different color to account for litter variability. Note that *Zmym2^-/-^* mice show reduced somite count within each litter. p-value calculated on basis of one-tailed t-test. **D.** Images of E8.5 *Zmym2^+/+^* and *Zmym2^-/-^* mice. **E.** Quantitative reverse-transcription PCR (qRT-PCR) of four upregulated genes expressed from promoter *(Ddx4, Tdrd1, Sohlh2, Hormad1)* and LINE-fusions *(Gc, Reg2)*. Each dot reflects the ratio of expression in KO/WT littermate embryos, with expression normalized to *RPS16*.

**Figure S2.**
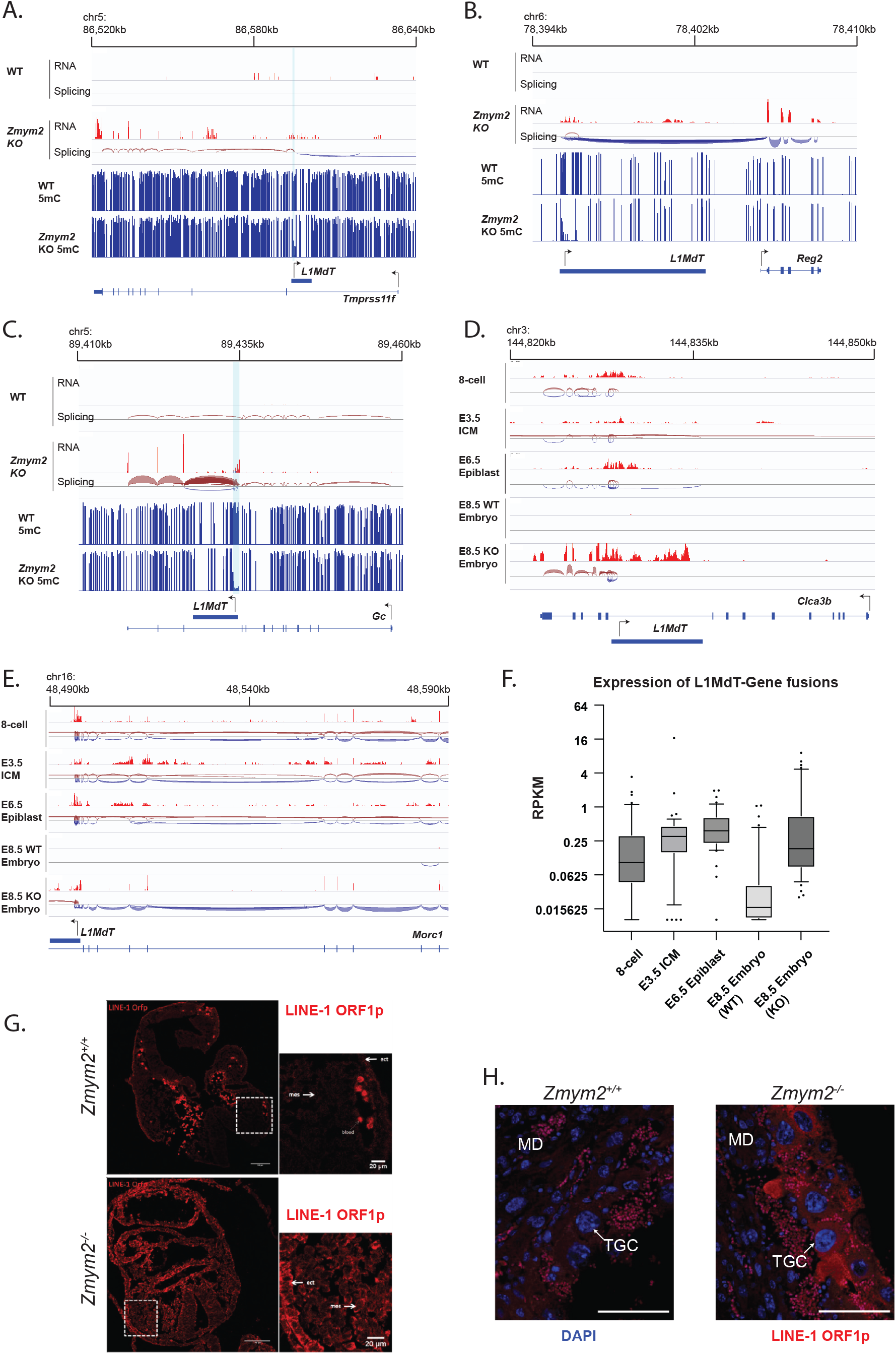
Expression of L1MdT-gene fusion transcripts in *Zmym2^-/-^* embryos and early embryonic development. **A-C.** Representative examples of L1MdT-gene fusion transcripts. Note examples of bidirectional transcription of an internal L1MdT element (**A**), splicing from an upstream L1MdT in a forward direction into a gene (**B**) and splicing from an internal L1MdT promoter in a forward direction into a gene (**C**). Hypomethylated DMRs are indicated in light blue. **D, E** Representative examples of L1MdT-gene fusion transcript expression in early embryonic development. **F.** Box plot of L1MdT-gene fusion transcripts during early mouse embryonic development. In **D-F,** 8-cell, E3.5 and E6.5 RNA-seq data are from Smith 2017. **G.** Immunofluorescence staining of LINE-1 ORF1p (red) in E8.5 *Zmym2^+/+^* and *Zmym2^-/-^* embryos. Regions of mesoderm and ectoderm in inset are indicated. Scale bar= 20μm **H.** Immunofluorescence staining of LINE-1 ORF1p (red) and DAPI (blue) of E9.5 *Zmym2^+/+^* and *Zmym2^-/-^* implantation sites. TGC= trophoblast giant cell, MD= maternal decidua. Small red cells visible in both samples are maternal blood cells. Scale bars= 100μm.

**Figure S3.**
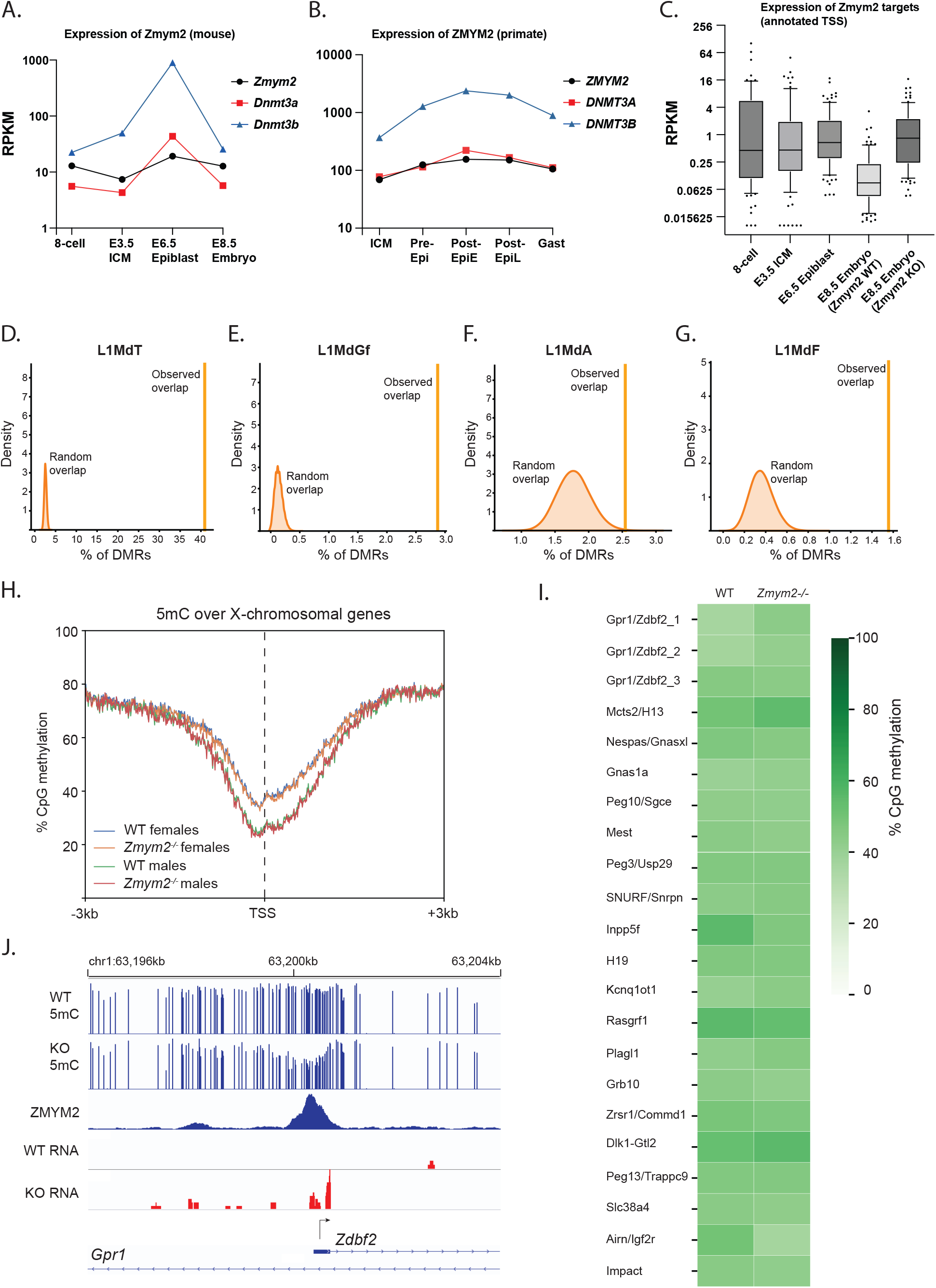
DNA methylation patterns in *Zmym2^-/-^* embryos. **A.** Expression of *Zmym2* and the *de novo* methyltransferases *Dnmt3a* and *Dnmt3b* during early murine embryonic development. Data from Smith 2017 (8 cell – E6.5) and this paper (E8.5). **B.** Expression of *ZMYM2* and the *de novo* methyltransferases *DNMT3A* and *DNMT3B* during cynomolgus monkey development. Data from Nakamura 2016. **C.** Expression of the 95 upregulated genes expressed from TSS during early murine embryonic development. Data from Smith 2017 (8 cell – E6.5) and this paper (E8.5). **D-G.** Monte-Carlo simulation was undertaken in which the positions of DMRs were randomly placed in the genome and repeated 100,000 times. The overlap with indicated transposon class is indicated as “Random overlap”. This number is contrasted with the “observed overlap” number for actual DMRs. Results are shown for L1MdT (**D)**, L1MdGf (**E)**, L1MdA (**F**), L1MdF (**G)**. **H.** Metaplot of DNA methylation over X-chromosomal genes in *Zmym2^+/+^* and *Zmym2^-/-^*. Note higher methylation in female than male, but no difference between *Zmym2^+/+^* and *Zmym2^-/-^*. **I.** DNA methylation over murine imprinted regions in *Zmym2^+/+^* and *Zmym2^-/-^*. No abnormalities are apparent. **J.** Hypomethylation, and aberrant transcription from the *Liz/Zdbf2* transient imprint in *Zmym2^-/-^*. Note enrichment of ZMYM2 at the target locus.

**Figure S4.**
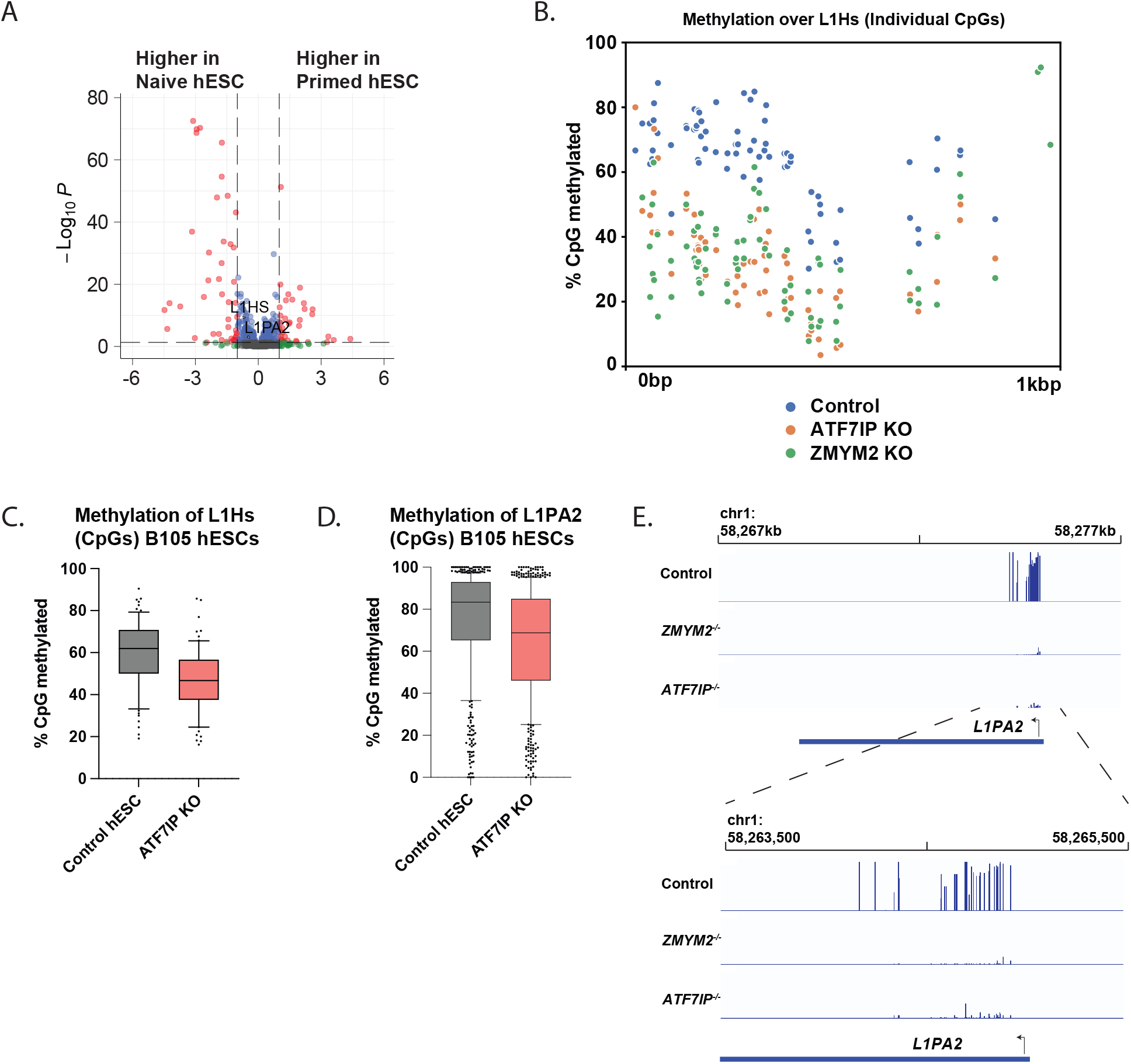
Upregulation and hypomethylation of young LINE elements in *ZMYM2^-/-^* human cells. **A.** Relative expression of transposon families in naïve and primed hESCs. Note that naïve hESCs show only a very modest increase in L1Hs and L1PA2 levels. **B.** Methylation levels of individual CpGs in an L1Hs consensus element for the genotype indicated. Only CpGs with >10-fold coverage in the Bar 2021 RRBS data are shown. **C.,D.** DNA methylation of L1Hs (**C**) and L1PA2 (**D**) control and *ATF7IP^-/-^* hESCs in B105 hESCs line. Note similarity to Figure 4B,C in this second hESC line. **E.** Example of an L1PA2 element hypomethylated in *ZMYM2^-/-^* and *ATF7IP^-/-^*. Inset is shown beneath main figure.

**Figure S5.**
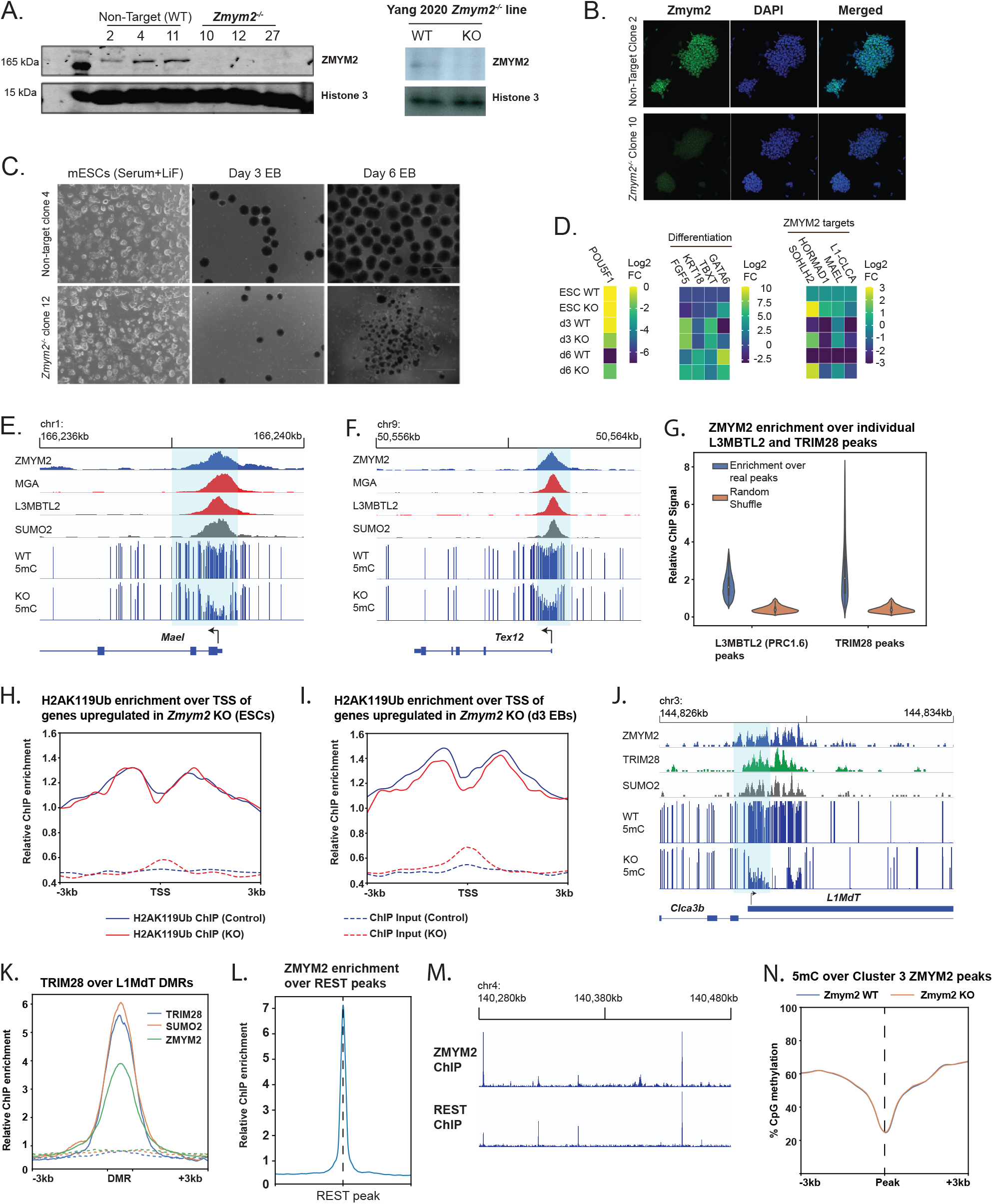
ZMYM2 targeting to PRC1.6, TRIM28 and REST binding sites. **A.** Western blots of control and *Zmym2^-/-^* clonal mESC lines generated on a V6.5 background (left) as well as WT J1 and *Zmym2^-/-^* mESCs generated by Yang and colleagues. **B.** Immunofluorescence staining of control and *Zmym2^-/-^* clonal line for ZMYM2. **C.** Light microscopy images of *Zmym2^+/+^* and *Zmym2^-/-^* ESCs and embryoid bodies. **D.** Relative expression of *POU5F1*, differentiation genes, and ZMYM2-target genes in control J1 mESCs and *Zmym2^-/-^* mESCs generated by Yang and colleagues, during embryoid body differentiation. **E., F.** ChIP-seq and DNA methylation data over the *Mael* **(E)** and *Tex12* **(F)** loci. Note colocalization of ZMYM2 with PRC1.6 components and hypomethylation in *Zmym2^-/-^*. Hypomethylated DMRs are indicated with light blue boxes. **G.** ZMYM2 ChIP-seq enrichment over L3MBTL2 and TRIM28 peak sets. Enrichment over random peaks of the same size is shown as a control. **H, I.** Relative ChIP enrichment of H2AK119Ub in *Zmym2^+/+^* and *Zmym2^-/-^* mESCs (**H**) and day 3 EBs (**I**) over genes upregulated in *Zmym2^-/-^* (transposon-fusion genes excluded). Note that there is no difference between *Zmym2^+/+^* and *Zmym2^-/-^*. The modestly increased input enrichment in *Zmym2^-/-^* may reflect increased locus accessibility as a result of impaired silencing. **J.** ZMYM2, TRIM28, SUMO2 ChIP-seq and WGBS data plotted over the L1MdT-*Clca3b* fusion transcript. Note correspondence of the three proteins at L1MdT promoters and corresponding reduction in DNA methylation. **K.** Metaplot of TRIM28, SUMO2 and ZMYM2 mESC ChIP-seq signal over DMRs hypomethylated in *Zmym2^-/-^* which overlap with L1MdT elements. Dashed lines indicate ChIP input. **L.** ZMYM2 ChIP-seq enrichment over REST binding sites. **M.** Overlap of REST and ZMYM2 binding sites over a stretch of chromosome 4. **N.** DNA methylation over Cluster 3 (REST-enriched) peaks in *Zmym2^+/+^* and *Zmym2^-/-^* embryos.

**Figure S6.**
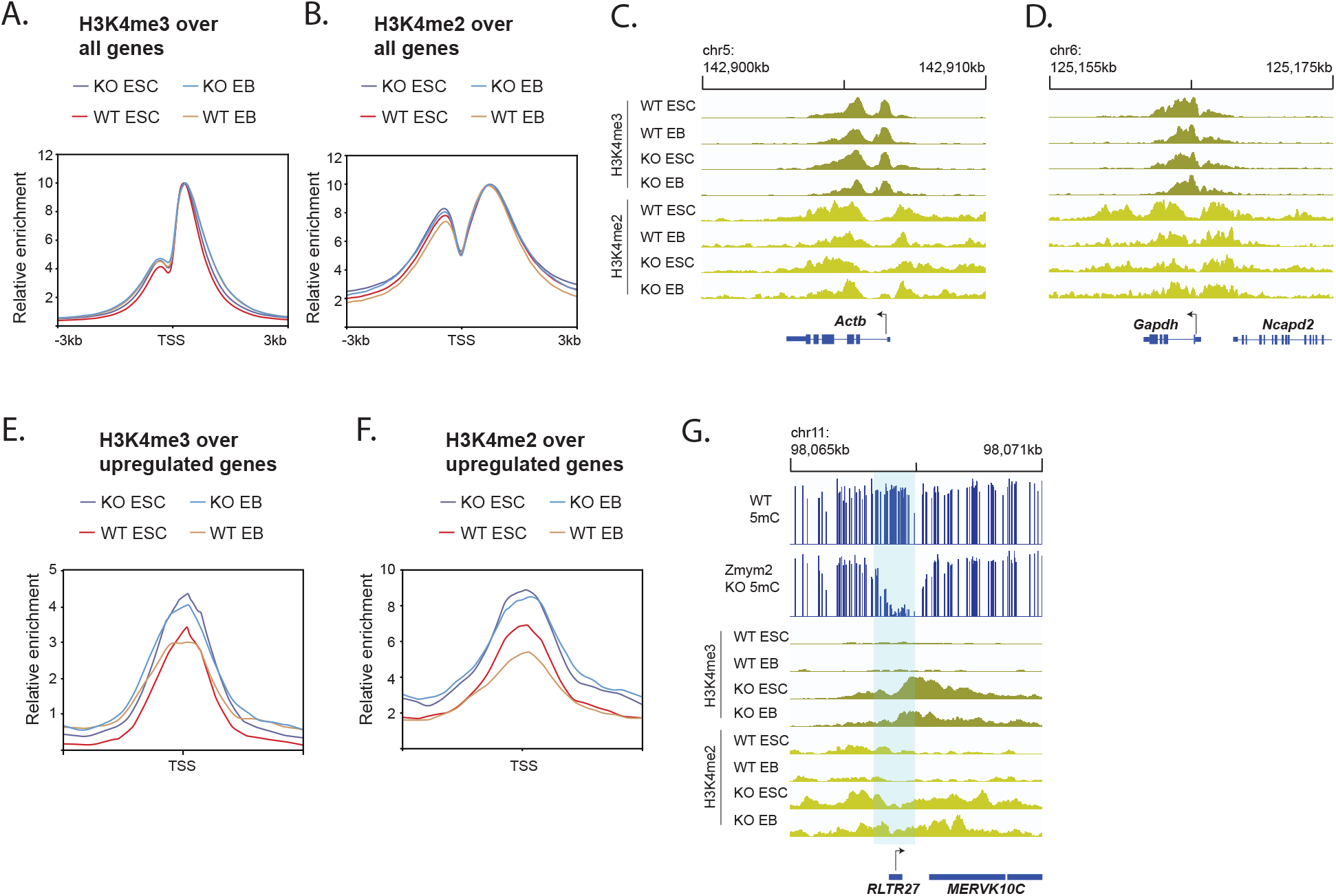
Aberrant H3K4 hypermethylation at ZMYM2-target sites in *Zmym2^-/-^* mESCs. **A., B.** Metaplot of H3K4me3 (**A**) and H3K4me2 (**B**) over all genes after normalization in indicated samples. Note similar pattern of enrichment over all replicates, including narrow H3K4me3 and broader H3K4me2 enrichment. **C.,D.** H3K4me3 and H3K4me2 ChIP enrichment over housekeeping *Actb* and *Gapdh* genes. **E, F.** H3K4me3 and H3K4me2 ChIP-seq enrichment over genes upregulated in *Zmym2^-/-^* embryos. **G.** A striking example of increased H3K4me2/3 over a hypomethylated DMR, an RLTR27 transposon in this case, in *Zmym2^-/-^* mESCs.

**Table S1. Mapping statistics for next generation sequencing data generated or used in the course of this project.**

**Table S2. RPKMs of all genes in control and *Zmym2^-/-^* mice.**

**Table S3. Sequences of primers used in this project.**

**Table S4. Annotations of upregulated genes in *Zmym2^-/-^* mice.**

**Table S5. Locations of DMRs in *Zmym2^-/-^* mice.**

## MATERIALS & METHODS

### Mice

Parental *Zmym2^+/-^* mice were generated by the Transgenic Core Facility of the Goodman Cancer Institute in a C57BL/6 background using a CRISPR-Cas9 targeting approach as described^15^. Timed mating was performed, and presence of a copulatory plug was identified at E0.5. Animals and experiments were kept in accordance with the standards of the animal ethics committee of McGill University, and the guidelines of the Canadian Council of Animal Care.

### Embryo collection and processing

All embryos were dissected in cold PBS and fixed for 20 minutes in 4% paraformaldehyde at 25°C. Following imaging of embryos with a Zeiss Lumar V12 stereomicroscope, samples used for tissue analysis were processed for either cryosection or paraffin embedding. Cryo-embedded samples were flash frozen in O.C.T. compound and sectioned to obtain 8- to 10-μm thick sections as described^62^. Paraffin-embedded samples were processed and embedded by the GCI Histology Core Facility and 6μm serial sections were obtained.

### Immunofluorescence staining of tissue sections

Immunofluorescence analyses were performed as described^63^. Anti-LINE-1 ORFp (Abcam cat#ab216324) was used at 1:100 dilution in PBS.

### Embryonic Stem Cell Culture Conditions

Both wild-type control and *Zmym2^-/-^* mESCs were generated from V6.5 ESCS (Novus Biologicals NBP1-41162) derived from C57Bl/6 x 129/sv cross. mESCs were cultured on 0.1% gelatin (EmbryoMax, ES-006-B) coated plated and media was refreshed every day. Cells were passaged every 2-3 days using 0.05% Trypsin-EDTA (Gibco, 25300062). These cells were cultured in Knockout™ D-MEM (Gibco, 10829018) media supplemented with 1000U/mL ESGRO Recombinant Mouse LIF (Millipore Sigma, ESG1107), 2 mM GlutaMAX (Gibco, 35050061), 100 μg/mL Primocin (InvivoGen, ant-pm-2), 0.1 mM 2-mercaptoethanol and 15% ES-qualified fetal bovine serum (Gibco, 10439024). All mESCs were mycoplasma-free and were periodically tested for mycoplasma contamination.

### Immunofluorescence staining of mESCs

mESCs were cultured on glass coverslips coated with 0.1% gelatin. Cells were then fixed with 4% paraformaldehyde for 15 minutes at room temperature and washed twice with 1xPBS. The mESCs were then permeabilized and blocked using 0.1% Triton-X diluted in 5% solution of donkey serum for 15 minutes at room temperature and washed twice with 1xPBS-T (0.1% Tween20 in 1xPBS). Primary antibody (ZMYM2, Invitrogen PA5-83208 at 1:1000 dilution) was added to the cells and incubated overnight at 4 °C. After overnight incubation, cells were washed twice with 1xPBS-T and secondary antibody and DAPI fluorophores were added (Donkey anti-Rabbit Alexa Fluor™ 488, A-21206 as a 1:500 dilution) for 45 minutes at room temperature. Cells were then washed twice with PBS-T and coverslips were mounted on microscopy slides using ProLong Gold. Cells were imaged on the LSM710 confocal microscope.

### RNA Isolation, Quantitative PCR and RNA sequencing of embryos

qPCR: Total mRNA was extracted from dissected embryos using a RNeasy micro kit (Qiagen-CAT# 74084) or AllPrep DNA/RNA Micro Kit (Qiagen-CAT# 82084). mRNA was reverse transcribed with MMLV (Invitrogen) according to manufacturer’s procedures. Real-time quantitative PCR was performed using TransStart^®^ Tip Green qPCR SuperMix (AQ141-01) on Realplex2 Mastercycler (Eppendorf).

RNA sequencing: Total RNA was isolated and sequenced from whole E8.5 embryos. Sequencing libraries were prepared by Genome Quebec Innovation Centre (Montreal, Canada), using the NEB mRNA stranded Library preparation. cDNA libraries were sequenced using the Illumina NovaSeq 6000 S2 sequencer, 100 nucleotide paired-end reads, generating 40-75 million reads per sample.

### RNA-sequencing analysis

RNA-seq fastq reads were trimmed using Trimmomatic(v0.34) to remove low-quality bases and remove adapters. Filtered reads were then aligned to the mm10 reference genome using STAR(v2.7.8a) with default parameters to generate .bam files. Picard(v2.9.0) was then used to sort and mark duplicates from the aligned bam files. Raw and normalized reads were the quantified using HTSeq-count and the StringTie suite(v1.3.5). The raw read counts were then used for differential expression analysis using DESeq2 and genes showing a minimum fold change of 4 and FDR(q-value) of less than 0.05 were selected as significant differentially expressed genes. Gene ontology over-representation was determined using clusterProfiler.

The TEtranscript pipeline was used to determine differential expression of transposons classes in Zmym2^-/-^ E8.5 embryos. RNA-seq reads were first trimmed using Trim Galore! (v0.6.6) with default parameters. Trimmed reads were then mapped to the mm10 genome using STAR aligner supporting multi-alignments per read (--winAnchorMultimapNmax 200, -- outFilterMultimapNmax 100 and a curated GTF file from the TEtranscript website). The resulting BAM files were used as inputs for the TEtranscript pipeline using default parameters and curated annotation refGene and repeatMasker files from the TEtranscript website. DESeq2 was then used to calculate differentially expressed transposon classes in *Zmym2^-/-^* embryos.

### Genetic Ablation of Zmym2 by CRISPR-Cas9 Nucleofections

*Zmym2^-/-^* mESCs on a V6.5 background were generated using the same sgRNA sequence used to generate the *Zmym2^-^* murine allele: AATGTTACAACCTTAGAAAC. CRISPR-Cas9 and sgRNA were delivered using the Lonza Biosciences 4D-Nucleofector to electroporate ribonuclear particles within cells. 300K cells were diluted in P3 primary cell solution prior to nucleofection and were hastily passaged to pre-warmed serum-LIF media after nucleofection. Clonal lines were generated by picking and expanding individual mESC colonies. Clonal mESC lines were then validated using PCR and Sanger Sequencing, western blotting and immunofluorescence. Control lines were generated by nucleofection with a non-targeting sgRNA and otherwise treated identically to *Zmym2^-/-^* lines *Zmym2^-/-^* mESCs in a CCE background were donated by the lab of Jianlong Wang^16^.

### Reverse-transcriptase quantitative PCR

Total mRNA was isolated from both wild-type, non-target and *Zmym2^-/-^* mESCs using RNAzol RT (Molecular Research Center Inc, RN 190) and first-strand cDNA was generated using the SensiFast cDNA Synthesis kit (FroggaBio, DD-BIO-65053). Quantitative PCR was then performed using PowerUP SYBR^™^ Green PCR (Invitrogen, A25742) on the Quantstudio 5 (Applied Biosystems) with the following cycling conditions: 50°C 2 minutes, 95°C 20 seconds, 45x (95°C 3 seconds, 60°C 30 seconds), 95°C 1 second). The qPCR reaction was performed using 1x concentration of PowerUP SYBR Green Master Mix, cDNA corresponding to 5ng of template mRNA and 0.5 μM of primer mix in a total of 6 μL reaction. The expression of ZMYM2 target genes were normalized to the housekeeping gene GAPDH and the sequence of primer used for qPCR are provided in Table S3.

### Western Blot

Protein was extracted using ice cold 1xRIPA lysis buffer supplemented with fresh protease inhibitors (1mM phenylmethylsulfonyl fluoride, 10mM sodium fluoride and 1mM sodium orthovanadate). Cell pellets were the subjected to 5 cycles of freeze-thawing using liquid nitrogen to ensure complete breakage of both the cell and nuclear membranes. Lysate protein concentration was measured using a Bradford Assay and 30-40μg of protein was run on a 6%-12.5% gradient SDS-PAGE. The resolved proteins were transferred to a polyvinylidene fluoride (PVDF) membrane, which was blocked using 5 mL of LI-COR Odyssey Blocking Buffer for 1h at room temperature. The membrane was then incubated overnight with primary antibody in 1xOdyssey Blocking Buffer 0.15% Tween-20 at 4°C. anti-ZMYM2 (ThermoFisher PA5-83208) was used at a 1:1000 dilution and anti-H3 (Abcam ab1791) was used at a 1:10,000 dilution. The membrane was then washed three x 5 minutes in PBS supplemented 0.1% Tween-20 and then incubated with secondary antibodies (LI-COR IRDye 680RD, 1:20,000 dilution) diluted in Odyssey Blocking Buffer and 0.15% Tween-20 for 1 h at room temperature. Membranes were then washed for 5 minutes twice in PBS 0.1% Tween and kept in PBS before imaging on the LI-COR imaging system.

### Whole-Genome Bisulfite Library Preparation

WGBS libraries were prepared from 500 ng of genomic DNA. Isolated genomic DNA (500 ng) of E8.5 mouse embryos (QIAamp DNA mini kit, 51304 or AllPrep DNA/RNA Micro Kit, Qiagen, 82084) and spike-in lambda DNA (1.25 ng) (New England Biolabs) was sheared using the M220 ultrasound sonicator (Covaris) to an average genomic size of 350 bp. 200 ng of sheared DNA was bisulfite-converted using the EZ DNA Methylation-Gold Kit (Zymo Research, D5005) using manufacturer’s instructions. Up to 100 ng of genomic DNA was then used to generate WGBS libraries using the Accel-NGS Methyl-Seq DNA Library prep kit (Swift Biosciences, 30024) according to manufacturer’s instructions and 13 cycles of PCR for final library amplification. Final concentration of libraries were determined using the Qubit 1X dsDNA High Sensitivity Assay Kit (Invitrogen, Q33230) and final library sizes were assessed by agarose gel electrophoresis. Libraries were then sequenced in paired-end using the NovaSeq 6000 at the Center for Applied Genomics operated by the SickKids Research Institute.

### Whole-Genome Bisulfite Sequencing Analysis

WGBS reads were trimmed to remove low-quality bases and the first five bases of R1 and ten bases of R2 were removed using Trim Galore! v(0.6.6) software (parameters: -q 20, --clip_R1 5, --clip_R2 10). Reads were then aligned to the mm10 reference genome using Bismark (v0.22.3) with default parameters. Aligned reads were then deduplicated and filtered for incomplete bisulfite conversion. Methylation calling over cytosines was done using Bismark and overall DNA methylation was calculated as a mean of all cytosine bases in CpG context. Metaplots analysis of CpG methylation over gene bodies or promoter regions were generated by calculating the percentage of CpG methylation of each RefSeq gene and 3Kb flanking regions. Differentially methylated regions were identified using the methylKit R package of 1000 bp tiled regions with a minimum of 10 read coverage, a difference of >25% methylation and a q-value of < 0.05.

### ChIP of Histone Modifications

Both wild-type and *Zmym2^-/-^* mESCs were crosslinked in 1% paraformaldehyde for 10 minutes and quenched with the addition of 1 mM of Glycine for 10 minutes to stop crosslinking. Cells were then lysed using lysis buffer solutions and sonicated with the M220 ultrasonicator (Covaris) in 1 mL tube with an AFA Fiber and the following conditions (cycles/burst = 200, Duty Factor = 20%, Peak Intensity Power = 75, time = 10 minutes and Temperature = 7 °C). The sheared DNA was then pre-cleared using magnetic beads (Sera-Mag Protein A/G SpeedBeads, VWR 17152104010150) for 1 hour to remove non-specific binding. Using a magnetic rack, the sonicated lysate was then separated to a new tube and primary antibodies was added (anti-H2AK119Ub Cell Signaling D27C, anti-H3K4me3 Cell Signaling CD42D8, anti-H3K4me2 Cell Signaling C64G9) and incubated overnight. Magnetic beads were then washed and added to the sonicated lysate and incubated for 2 hours at 4 °C to allow for the beads to bind to the antibody/protein/DNA complex. Beads were then washed with buffer of increasing salt concentration (Wash Buffer A: 50mM HEPES, 1% TritonX-100, 0.1% Deoxycholate, 1mM EDTA, 140 mM NaCl and Wash Buffer B: 50 mM HEPES, 0.1% SDS, 1% TritonX-100, 0.1% Deoxycholate, 1 mM EDTA, 500 mM NaCl) and TE buffer. Protein/DNA complexes of interest were finally eluted using elution buffer (50 mM Tris-HCl, 1 mM EDTA, 1% SDS) incubated at 65 °C for 10minutes and separated from the beads using a magnetic rack. Samples were then decrosslinked by incubating overnight at 65 °C. Residual RNA and protein were removed with the addition of RNAseA and proteinase K. DNA was then purified using the Qiagen Minelute PCR Purification Kit using manufacturer’s instructions.

### CUT&RUN of L3MBTL2

To determine the localization of L3MBTL2 in Zmym2^+/+^and Zmym2^-/-^ mESCs, CUT&RUN was used using manufacturer’s instructions (EpiCypher CUTANA ChIC/CUT&RUN Kit, 14-1048). The eluted DNA from the CUT&RUN or ChIP was kit was then used to construct multiplex libraries with the NEBNext Ultra II DNA library Kit using manufacturer’s instructions.

### ChIP-seq and CUT&RUN Data analysis

Reads were trimmed using Trimmomatic (v0.6.6) using default parameters and then aligned to the mm10 genome using BWA(v0.7.17). Removal of PCR duplicates was then done using Picard (v2.0.1). MACS2 was then used for peak identification using default parameters and inputs as controls. Aligned BAM files were used to generate the bigwigs using DeepTools. The resulting bigwigs were used for metaplot and heatmaps using the computeMatrix and plotHeatmap functions in DeepTools.

## ACKNOWLEDGEMENTS

We thank the Genome Quebec Innovation Centre, the Centre for Applied Genomics at SickKids Hospital, the McGill University Advanced BioImaging Facility (ABIF), and the Rosalind and Morris Goodman Cancer Institute Histology Innovation Platform for assistance. We thank Mitra Cowan and the McGill Integrated Core for Animal Modeling for assistance generating and maintaining mice. We thank Jianlong Wang for donating *Zmym2^-/-^* mESCs. The research was funded by CIHR Project Grant PJT-159768 (M.B), CIHR Project Grant PJT-166169 (W.P.), the New Frontiers in Research Fund Grant NFRFE-2018–00883 (W.P.) and the NSERC Grant RGPIN-2018-04856 (W.P.). A.G. was supported by a CIHR Canada Graduate Scholarships Doctoral Award. Q.Z. was supported by doctoral training awards by the FRQS and JST. G.B. was supported by a Canada Research Chair and an FRQS Chercheurs-mérite. W.P. was supported by an FRQS Chercheurs-boursier.

## AUTHOR CONTRIBUTIONS

A.G., D.S., J.S., and M.K. conducted experiments. D.S., I.H., Q.Z. and S.K. conducted bioinformatic analysis. Y.Y. provided guidance on phenotypic analysis of mice. G.B. provided guidance on bioinformatic analysis. M.B. and W.P. supervised the research project. A.G., D.S. and W.P. wrote the manuscript.

